# Progression from feature-specific brain activity to hippocampal binding during episodic encoding

**DOI:** 10.1101/735761

**Authors:** Rose A. Cooper, Maureen Ritchey

**Affiliations:** Department of Psychology, Boston College

## Abstract

The hallmark of episodic memory is recollecting multiple perceptual details tied to a specific spatial-temporal context. To remember an event, it is therefore necessary to integrate such details into a coherent representation during initial encoding. Here we tested how the brain encodes and binds multiple, distinct kinds of features in parallel, and how this process evolves over time during the event itself. We analyzed data from 27 human subjects (16 females, 11 males) who learned a series of objects uniquely associated with a color, a panoramic scene location, and an emotional sound while functional magnetic resonance imaging data were collected. By modeling how brain activity relates to memory for upcoming or just-viewed information, we were able to test how the neural signatures of individual features as well as the integrated event changed over the course of encoding. We observed a striking dissociation between early and late encoding processes: left inferior frontal and visuo-perceptual signals at the onset of an event tracked the amount of detail subsequently recalled and were dissociable based on distinct remembered features. In contrast, memory-related brain activity shifted to the left hippocampus toward the end of an event, which was particularly sensitive to binding item color and sound associations with spatial information. These results provide evidence of early, simultaneous feature-specific neural responses during episodic encoding that predict later remembering and suggest that the hippocampus integrates these features into a coherent experience at an event transition.

**SIGNIFICANCE STATEMENT:** Understanding and remembering complex experiences is crucial for many socio-cognitive abilities, including being able to navigate our environment, predict the future, and share experiences with others. Probing the neural mechanisms by which features become bound into meaningful episodes is a vital part of understanding how we view and reconstruct the rich detail of our environment. By testing memory for multimodal events, our findings show a functional dissociation between early encoding processes that engage lateral frontal and sensory regions to successfully encode event features, and later encoding processes that recruit hippocampus to bind these features together. These results highlight the importance of considering the temporal dynamics of encoding processes supporting multimodal event representations.

## INTRODUCTION

Our ongoing perceptual experience of the world includes a stream of disparate, multimodal features unfolding in parallel. Memory-related increases in brain activity during encoding are often found in visuo-perceptual brain regions (Spaniol et al., 2009; Kim, 2011), emphasizing that stronger, more precise representations of perceptual information support memory formation. For instance, comparing encoding of different kinds of information, such as words and pictures, reveals category-selective patterns of sensory activity that predict subsequent recollection of these individual features (Gottlieb et al., 2010; Duarte et al., 2011; Park and Rugg, 2011). Yet natural experience not only involves representing different perceptual features but, crucially, it requires us to encode them simultaneously. Moreover, we later remember this information as a unified event. In this study, we investigated how perceptual features are uniquely represented during encoding and the neural operations that bind them together.

Studies addressing these issues have largely examined memory for simple paired associations studied as short “events”. This research involves presenting subjects with pairs of features within the same trial, such as a location, person, emotion, or color. By predicting subsequent memory separately for each component feature (Uncapher et al., 2006; Staresina and Davachi, 2008; Gottlieb et al., 2012; Ritchey et al., 2018) or contrasting events with overlapping or non-overlapping feature types (Horner et al., 2015), these studies take a step toward understanding how the brain represents and encodes distinct kinds of information in parallel. Moreover, some regions, most notably the hippocampus, show a preference for binding pairs of features during encoding rather than promoting memory for either detail alone (Uncapher et al., 2006; Horner et al., 2015). However, restricting encoding trials to bimodal associations leaves it unclear how distinct feature signals of more complex experiences are simultaneously distinguished, encoded, and integrated as the event unfolds.

The hippocampus is considered to be crucial for binding elements of our experience with contextual information (Davachi, 2006; Diana et al., 2007; Eichenbaum et al., 2007; Ranganath, 2010), though less is known about how exactly the hippocampus organizes multimodal information. Its involvement might be tied to total memory content, including details and their associations, during event encoding (Addis and McAndrews, 2006; Staresina and Davachi, 2008; Park and Rugg, 2011), such that activity tracks the amount of information subsequently recalled (Qin et al., 2011) regardless of its content or relational structure. Alternatively, the hippocampus could be specifically recruited to organize our environment in a structured way, perhaps around a spatial framework (Horner et al., 2015; Deuker et al., 2016; Zeidman and Maguire, 2016). It is also important to distinguish the role of the hippocampus in multimodal event encoding from that of other brain regions that also show encoding effects related to organization and integration, such as left inferior frontal gyrus (Ranganath et al., 2004; Addis and McAndrews, 2006; Staresina and Davachi, 2006; Park and Rugg, 2011).

Here, we present the encoding data from our prior work (Cooper and Ritchey, 2019) in which participants encoded and reconstructed complex events. In line with past research (Horner and Burgess, 2013; Joensen et al., 2019), we previously showed that successful recall of event associations in our task — an object with a color, scene location, and sound — exhibits a dependent structure (Cooper and Ritchey, 2019). In the current analyses, we leverage the multimodal aspect of this paradigm to test how features are prioritized and integrated during encoding. To preview the results, we found that visuo-perceptual and left inferior frontal regions supported feature-specific and binding processes, respectively. Surprisingly, however, hippocampal activity at the onset of events did not correlate with subsequent memory. Based on recent findings that hippocampal activity is particularly enhanced at the end of an event (Ben-Yakov et al., 2014; Ben-Yakov and Henson, 2018), we further investigated the temporal dynamics of event encoding. Specifically, we contrasted the neural processes supporting encoding of upcoming versus just-studied information, testing if hippocampal signals at an event transition act to bind episodic features in memory.

## MATERIALS AND METHODS

Portions of this dataset have been previously reported (Cooper and Ritchey, 2019). Whereas the previous paper was focused on functional connectivity analyses of the retrieval phase, the current paper reports on univariate activation analyses of the encoding phase. Methods for MRI data collection, the task procedure, and behavioral analyses have been previously detailed in (Cooper and Ritchey, 2019), and so are summarized here.

### Participants

27 participants took part in the current experiment (16 females, 11 males). All participants were 18-35 years of age (mean = 21.7 years, SD = 3.58) and did not have a history of any psychiatric or neurological disorders. Seven additional subjects took part but were excluded from data analyses: two participants did not complete the experiment, one due to anxiety and the other due to excessive movement in the MRI scanner, four additional participants had chance-level performance on the memory task, and one subject was excluded after data quality checks revealed 3/6 encoding functional runs exceeded our motion criteria. Informed consent was obtained from all participants prior to the experiment and participants were reimbursed for their time. Procedures were approved by the Boston College Institutional Review Board.

### Experimental Design and Statistical Analyses

#### Paradigm

Participants were presented with a series of 144 unique object “events” in an MRI scanner, 24 per scan run. Each object was presented in a color from a continuous CIEL*A*B color spectrum, in a scene location within one of 6 panoramic environments, and in conjunction with one of 12 sounds - 6 that were emotionally negative and 6 that were neutral. All sounds contained natural, easily recognizable content and were 6 seconds in duration, corresponding to the time each event was displayed during encoding. Events were separated by a 1 second fixation. Participants were instructed to integrate the object and its associated features into a meaningful event, but no response was required (Figure 1A). Allocations of features to objects as well as the presentation order of events were randomized within each subject. This memory encoding phase is the focus of all fMRI analyses presented here; see Cooper and Ritchey (2019) for results from the retrieval phase. After encoding 24 events, subjects were tested on their memory for the features associated with each object, presented in grayscale as memory cues. On each retrieval trial, participants attempted to remember all of the features in their mind for 4s, following which they were prompted to remember if the object was paired with a negative or neutral sound (2s), and to reconstruct the object’s color and spatial location (6s each). For the sound feature, participants reported their confidence in their decision (‘maybe’ or ‘sure’), whereas, for the visual features, participants changed the object’s color and moved it around the panorama to recreate its visual appearance as precisely as possible in 360-degree space.

**Fig.1.**
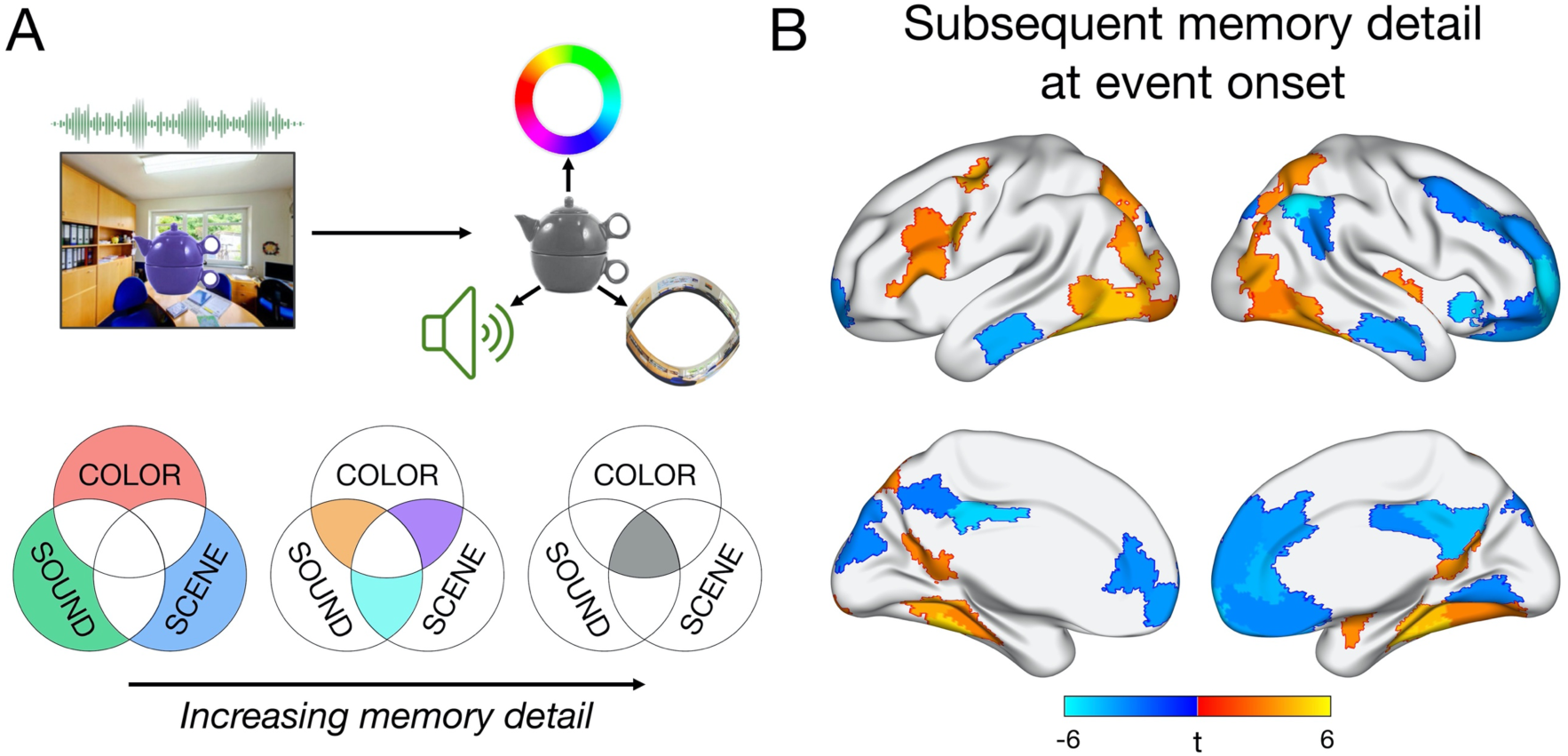
Task design and neural correlates of subsequent memory detail at the onset of each multi-feature event. A) During encoding, participants learned a series of events presented for 6s, each uniquely associating an object to a color, sound, and scene location. For subsequent memory analyses, each event was coded by its level of memory detail, specified by the number of features (0-3) that were successfully recollected. B) Significant relationships (*p* < .05 FDR corrected) between activity at event onset and the number of features subsequently recalled. All voxels within a functionally homogeneous region, defined based on a whole-brain parcellation, are color-coded by the region’s t-statistic.

#### fMRI Data Acquisition

MRI scanning was performed using a 3-T Siemens Prisma MRI scanner at the Harvard Center for Brain Science, with a 32-channel head coil. Structural MRI images were obtained using a T1-weighted (T1w) multiecho MPRAGE protocol (field of view = 256 mm, 1 mm isotropic voxels, 176 sagittal slices with interleaved acquisition, TR = 2530 ms, TE = 1.69/ 3.55/ 5.41/ 7.27 ms, flip angle = 7º, phase encoding: anterior-posterior, parallel imaging = GRAPPA, acceleration factor = 2). Functional images were acquired using a whole brain multiband echo-planar (EPI) sequence (field of view = 208 mm, 2 mm isotropic voxels, 69 slices with interleaved acquisition, TR = 1500 ms, TE = 28 ms, flip angle = 75º, phase encoding: anterior-posterior, parallel imaging = GRAPPA, acceleration factor = 2), for a total of 466 TRs encompassing encoding and retrieval trials per scan run. Fieldmap scans were acquired to correct the EPI images for signal distortion (TR = 314 ms, TE = 4.45/ 6.91 ms, flip angle = 55º).

#### Behavioral Data Processing

As previously reported (Cooper and Ritchey, 2019), participants’ responses for the object color and scene location questions were measured in terms of error — the difference between the target (encoded) feature value and the reported feature value in 360-degree space. Responses to the sound question were considered in terms of accuracy and confidence. Each encoding event was characterized in terms of its pattern of memory success (correct vs. incorrect subsequent retrieval) for each of these features. We *a priori* chose to use a more conservative method for identifying “correct” memory retrieval than in our previous analysis of this data (Cooper and Ritchey, 2019) to ensure that we only included trials associated with confident/precise subsequent recollection, and to more evenly balance the distribution of trials across different remembered feature combinations. Specifically, the sound feature was considered correctly recalled if the participant successfully identified the associated sound as negative or neutral *and* reported this memory with high confidence. For color and scene location features, we first fitted a mixture model (Zhang and Luck, 2008; Bays et al., 2009) to the aggregate group error data, per feature. The mixture model includes a uniform distribution to estimate the proportion of responses that reflect guessing, as well as a circular Gaussian (von Mises) distribution to estimate the proportion of responses that reflect successful remembering. We defined “correct” color and scene memory by the probability that an error had at least a 75% chance of belonging to the von Mises distribution and not the uniform distribution, resulting in a threshold of +/- 42 degrees for color and +/- 24 degrees for scene memory. Whereas participant-level models could capture variation in memory precision between subjects (with participant-specific thresholds for ‘correct’/’incorrect ‘memory), we chose to define memory based on group-level data. Modeling at the subject level may produce variable threshold estimates because of differences in model fit rather than differences in memory per se. Moreover, trials allocated as “correct” or “incorrect” across subjects would not be directly comparable. In line with our prior work (Richter et al., 2016; Cooper et al., 2017; Cooper and Ritchey, 2019), we felt that a threshold for correct memory based on group data was therefore most appropriate.

For each encoding event, a composite measure of memory detail was calculated as the number of features subsequently recalled (0-3). Of note, this is a slightly different composite measure of memory than we have used previously (Cooper and Ritchey, 2019), where we accounted for variations in confidence and precision within “successful” recall. However, we have already restricted our measure of successful memory for each feature to a confident and precise judgment, and, in our previous behavioral analyses, it was successful recall but not the precision of recall that showed a dependent structure in memory.

#### fMRI Data Preprocessing

All data preprocessing was performed using FMRIPrep version 1.0.3 (Esteban et al., 2018) with the default processing steps. Each T1w volume was corrected for intensity non-uniformity and skull-stripped. Spatial normalization to the ICBM 152 Nonlinear Asymmetrical template version 2009c was performed through nonlinear registration, using brain-extracted versions of both the T1w volume and template. All analyses reported here use images in MNI space. Brain tissue segmentation of cerebrospinal fluid (CSF), white-matter (WM) and gray-matter (GM) was performed on the brain-extracted T1w image. Functional data was slice time corrected, motion corrected, and corrected for field distortion. This was followed by co-registration to the corresponding T1w using boundary-based registration with 9 degrees of freedom. Six principal components of a combined CSF and WM signal were extracted using aCompCor (Behzadi et al., 2007) for use as nuisance regressors. Framewise displacement was also calculated for each functional run. For further details of the pipeline, including the software packages utilized by FMRIPrep for each preprocessing step, please refer to the online documentation: https://fmriprep.readthedocs.io/en/1.0.3/index.html. After preprocessing, the first 116 TRs were selected from each run, capturing all encoding trials and extending 6s beyond completion of the final trial. Encoding runs were checked for motion and were excluded from data analyses if more than 20% of TRs exceeded a framewise displacement of 0.3 mm. Participants were retained for analyses if at least 4/6 runs passed criteria. One subject was excluded as result of 3 runs having high motion, but no other subject had runs removed because of motion. Four participants successfully completed only 5 out of the 6 scan runs, 3 as a result of exiting the scanner early and 1 due to a technical problem with the sound system during the first run.

#### Whole Brain Parcellation

To test which regions across the whole brain were sensitive to subsequent memory effects, we divided the cortex into 200 functionally homogeneous regions using a recent cortical parcellation (Schaefer et al., 2018; https://github.com/ThomasYeoLab/CBIG). Four medial temporal areas were added to this whole-brain atlas (left and right hemisphere as separate regions). The hippocampus was obtained from a probabilistic medial temporal lobe (MTL) atlas (Ritchey et al., 2015; https://neurovault.org/collections/3731/) and amygdala was added from the Harvard-Oxford subcortical atlas, consistent with our prior methods (Cooper and Ritchey, 2019). We decided to implement a whole brain parcellation method rather than a more traditional voxel-based approach for two reasons: i) to utilize what we know about anatomical and functional similarities across the brain to maximize statistical power and ii) allow us to draw clearer conclusions about the function of individual brain regions defined in a discrete way. Although parcellation-based averaging has the potential to miss task-specific variations in activity within larger regions, we believe that using a fine-grained parcellation, with multiple-comparisons correction, may be an optimal approach for whole brain analyses that extend beyond hypothesis-driven functional regions of interest. The results of all whole brain analyses were displayed using BrainNet Viewer (Xia et al., 2013), where, for visualization purposes, every voxel within a region was allocated the same t statistic calculated from an analysis of that region’s mean activity.

#### Single Trial Analyses

All analyses were run using single trial estimates of activity during encoding. Single trial betas were computed using SPM12, implementing the least squares-separate (LS-S) method (Mumford et al., 2012) where a separate general linear model (GLM) was constructed for each encoding trial. In each single trial GLM, the first regressor corresponds to the trial of interest and a second regressor codes for all other encoding trials in that functional run. Functional data were denoised by including framewise displacement, 6 aCompCor principal components, and 6 movements parameters as nuisance regressors. Spike regressors were also included to exclude data points with high motion (> 0.6mm). Single trial models were constructed using: 1) a stick function at stimulus *onset*, or 2) a stick function at stimulus *offset*, with each type of regressor convolved with the canonical HRF. Mean single trial beta series were extracted by averaging across voxels within each of the 204 regions per subject. Analyses of single trial betas were conducted using RStudio (RStudio Team, 2016), R version 3.5.1.

Before running any analyses, data were first cleaned by z-scoring the single trial beta values within each region per subject and removing trials greater than 4 SD from the mean for any region. A total of 108 (2.85%) and 113 (2.98%) out of 3792 trials were removed for analyses of event onsets, used to test the relationship between activity and memory for the *upcoming* event, and offsets, used to test the relationship between activity and memory for the *preceding* event. All statistical tests quantify the relationship between each region’s encoding beta series and subsequent memory using linear mixed effects models (*lme4;* Bates et al., 2015), fitted using restricted maximum likelihood. In all models, subject was included as a random effect with a variable intercept and variable slopes for all fixed effects. This ‘maximal’ approach guards against an inflated Type 1 error rate (Barr et al., 2013) but can commonly result in model convergence failures with increasing complexity of the random effect variance-covariance structure (Matuschek et al., 2017). To address this problem, random effect correlations were constrained to zero in all models (Barr et al., 2013; Matuschek et al., 2017) using the formula *lmer(Y ~ X_1_ + X_2_ … + (1 + X_1_ + X_2_ … || Subject))*. To avoid singular fits for more complex models, the random effects structure of models with more than two fixed effects were additionally simplified using the *lmerTest* package (Kuznetsova et al., 2017) ‘step’ function, which iteratively removes random slopes that do not improve the model fit. Each fixed effect was tested against zero using *lmerTest*, which implements t-tests with Satterthwaite’s approximation method, estimating the effective degrees of freedom for each effect. All p-values reported were FDR-corrected (alpha = .05) for multiple comparisons across all 204 brain regions within each type of mixed effects model.

### Code Accessibility

Single trial encoding data for all subjects and brain regions as well as R scripts to run the analyses described here have been made freely available through GitHub: http://www.thememolab.org/paper-bindingfmri/. This repository also contains extended data in the form of csv and nifti files for the results of all whole-brain linear mixed effects analyses.

## RESULTS

### Neural correlates of subsequent episodic detail are sensitive to distinct memory features

We first sought to test where across the brain activity at the onset of a multi-feature event was sensitive to the amount of detail recalled (Figure 1B). Activity of each brain region was predicted from an objective measure of memory detail coding each trial according to whether 0, 1, 2, or 3 features were subsequently remembered. Significant memory-related increases in activity were found in a number of visuo-perceptual brain regions, including bilateral occipital and ventral visual regions, bilateral parahippocampal cortex and retrosplenial cortex, as well as right amygdala and superior temporal cortex. Left inferior frontal gyrus — frequently associated with successful associative encoding — also positively tracked the amount of episodic detail later recalled. In contrast, activity of default network regions, including bilateral medial prefrontal cortex (mPFC), posterior cingulate, and middle temporal gyrus, was negatively associated with increasing subsequent memory detail. Thus, left lateral frontal and visuo-perceptual regions appear to prioritize event features at the onset of encoding to support a detailed memory representation.

We next wanted to determine which of these regions were associated with subsequent memory in a feature-specific way. That is, to what extent does encoding a multi-feature event involve the parallel activation of feature-specific patterns? To this end, mixed effects models were used to identify patterns of event-onset activity that were *uniquely* predicted by subsequent memory for the individual features — color, sound, and scene perspective — with each feature predictor binary-coded according to whether memory was subsequently correct or incorrect. Of note, remembering any one feature in our paradigm involves an association of that feature to an object. Therefore, this analysis identifies regions sensitive to cued recall of different types of event content rather than recognition of single features.

Pronounced feature-specific effects emerged in terms of positive subsequent memory correlates (Figure 2). Specifically, successful encoding of the object’s color was associated with enhanced activity of right ventral visual cortical areas, whereas subsequent memory for the sound was positively related to activity in bilateral superior temporal (auditory) cortex. Neural correlates of successful scene encoding were more widespread, with positive changes in activity observed in bilateral dorsal medial parietal and occipital cortex, as well as bilateral retrosplenial and parahippocampal cortex, frequently reported to be sensitive to specific representations of spatial locations and perspectives (e.g., Epstein, 2008; Robertson et al., 2016; Robin et al., 2018; Berens et al., 2019). Thus, we were able to detect unique patterns of brain activity that positively tracked different kinds of episodic features encoded simultaneously, with the most widespread effects for encoding scene details. Notably, left inferior frontal gyrus, which tracked overall memory detail, did not significantly support memory for any individual feature. There were also signs of feature-specific negative subsequent memory effects (Figure 2), predominantly localized to the default network. Given evidence of distinct default network subsystems (Buckner and DiNicola, 2019), future work should consider the possible content-specificity of negative encoding effects.

**Fig.2.**
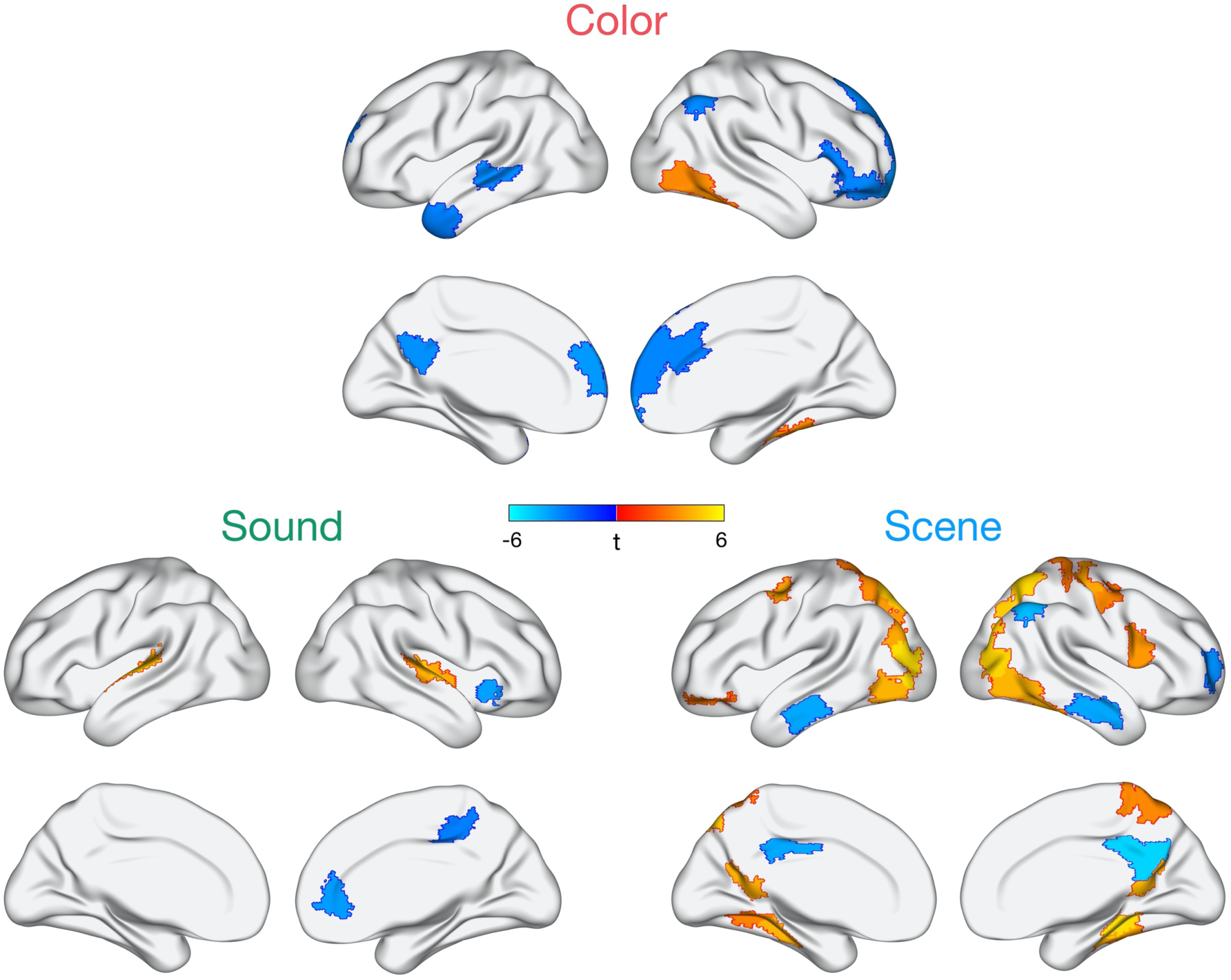
Regions whose activity at the onset of each event is uniquely predicted by subsequent color, sound, or scene memory while these features are encoded in parallel. Task-positive effects show mostly distinct patterns across the three features. Effects shown are FDR-corrected for multiple comparisons at *p* < .05, and all voxels within a region are color-coded by the region’s t-statistic.

### Memory-related neural activity shifts over the time course of encoding

The previous analysis showed that there are partially distinct, parallel patterns of encoding activity that support memory for individual features. Yet a fundamental aspect of episodic memory is the binding of multimodal features into a coherent event representation. While we found evidence that left lateral frontal cortex activity correlated with memory detail and thus may facilitate integration, surprisingly, we found that hippocampal activity was unrelated to the number of features recalled (|*betas*| < 0.02, |*t*s| < 0.26, *p*s_corrected_ > .91) or to memory for any feature alone (|*betas*| < 0.04, |*t*s| < 0.50, *p*s_corrected_ > .81). Hippocampal subsequent memory effects have been frequently (although not universally) reported in event-related paradigms (Kim, 2011), specifically for binding multiple episodic features (Horner et al., 2015), and most models of episodic memory assume that the hippocampus supports this binding process. In light of evidence that hippocampal activity may be particularly sensitive to the end of an event (Ben-Yakov et al., 2014; Ben-Yakov and Henson, 2018), we reasoned that some encoding-related brain regions might be important for prioritizing features early on (predicting memory for the upcoming event), whereas other regions, such as hippocampus, might be engaged later to integrate just-viewed information into a memory trace.

To test this hypothesis, we next examined whether the pattern of subsequent memory effects across the brain would shift over the course of encoding. Specifically, whereas the previous set of analyses were focused exclusively on how onset activity was related to memory for the upcoming event, we now considered how activity at the *offset* of each event (6s after onset) was related to subsequent memory for the just-viewed information (Figure 3A). Overall, fewer regions exhibited sensitivity to memory detail for the preceding event (Figure 3B), although the sensitivity of some default regions to memory detail was sustained over encoding, with an overlapping subset of mPFC regions showing a negative relationship between activity and subsequent memory for both upcoming and just-viewed event information. A transition of subsequent memory effects was particularly apparent for left inferior frontal and feature-selective regions — these regions did not significantly exhibit sensitivity to memory detail over the full course of encoding. Of particular interest, activity at event offset in left hippocampus was positively related to the number of event details subsequently remembered (left: *beta* = 0.14, *t* = 3.50, *p*_corrected_ = .024; right: *beta* = 0.11, *t* = 2.40, *p*_corrected_ = .099), reflecting the emergence of a hippocampal memory effect at the end of an encoding trial. This effect appeared to be relatively specific to the period after the event of interest, in that a control analysis modeling the entire trial as a 6-s block (onset to offset) did not reveal any significant hippocampal subsequent memory effects (|*betas*| < 0.03, |*t*s| < 1.72, *ps*_corrected_ > .19). Positive correlates of memory detail for the preceding event were also seen in left inferior parietal cortex, middle temporal gyrus, and temporal pole.

**Fig.3.**
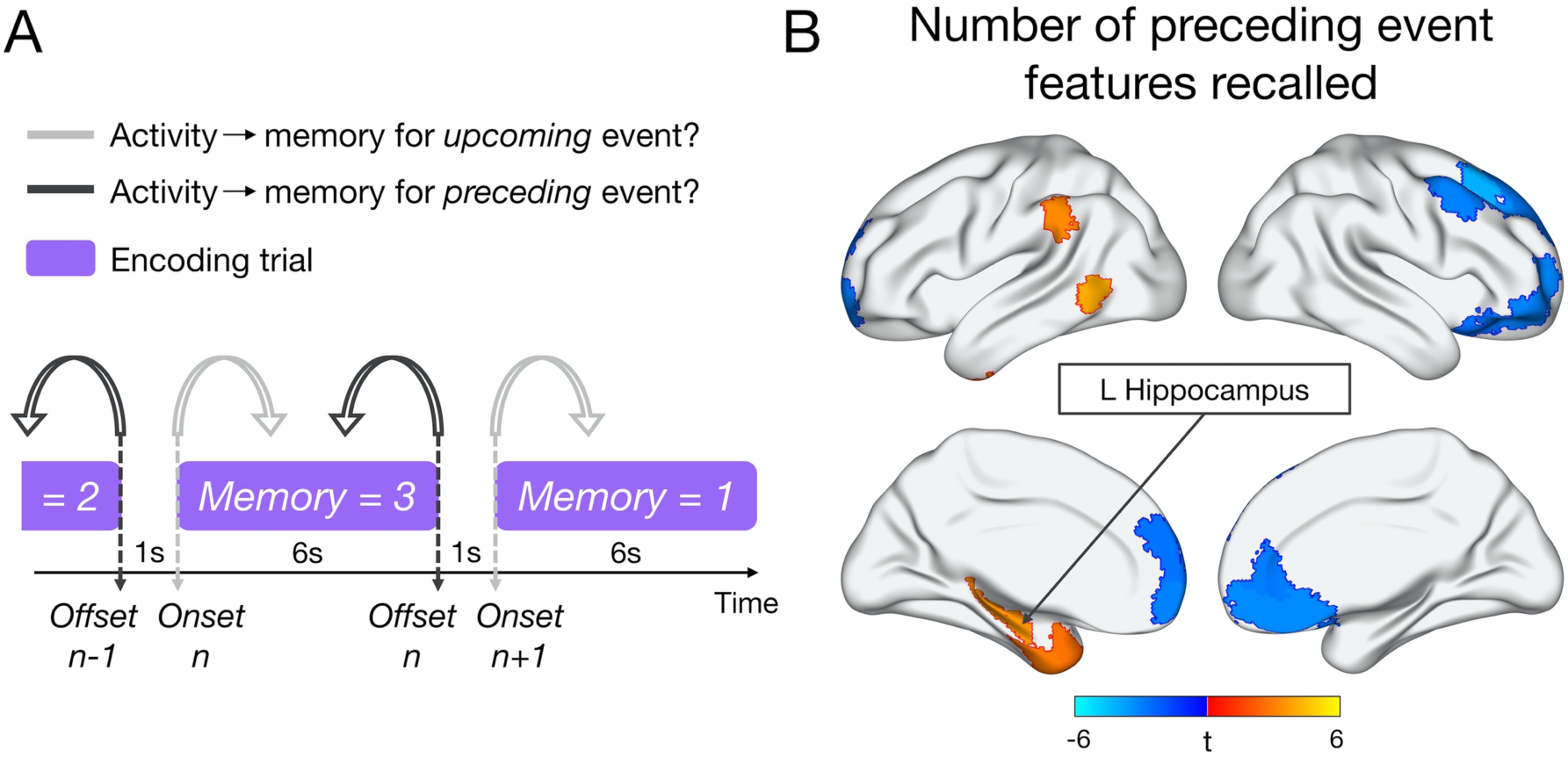
Analysis approach and neural correlates of preceding memory detail. A) Schematic of the different models relating brain activity at event onset (light gray) or offset (dark gray) to subsequent memory for the upcoming or preceding event, respectively. B) Brain regions whose activity at event offset is significantly (*p* < .05 FDR corrected) predicted by the number of details (0-3) subsequently recalled from the preceding event. All voxels within a region are color-coded by the region’s t-statistic.

Because this study was not originally designed with the intention of studying offset responses, it is important to consider the timing of event offsets relative to other trial components. Onsets and offsets for a given trial were placed 6s apart, allowing us to separately relate them to that trial’s memory score. In fact, within each ROI per subject, the mean correlation between activity at trial *N* onset and trial *N* offset (6s apart) was low (mean r = .14, SE = .01). In contrast, due to the fixed 1s ITI between trials, the offset and onset beta estimates of neighboring trials — offset *N* and onset *N+1* — were highly correlated (mean r = .88, SE = .02), as expected with a slow BOLD response. Importantly, however, these time points were associated with different memory scores (Figure 3), and control analyses show that memory scores for neighboring trials were largely independent of one another (total memory detail: mean r = .10, SE = .03; color: mean r = .01, SE = .02; sound: mean r = .11, SE = .04; scene location: mean r = .06, SE = .02). Moreover, to rule out the possibility of any low correlation between memory on adjacent trials influencing our results, we re-ran the analyses predicting onset *N* or offset *N* activity from memory detail on trial *N*, also including memory detail on the neighboring trial, *N-1* or *N+1*, respectively, as a covariate. The pattern of results did not change, with left inferior frontal gyrus tracking memory for the upcoming event (*betas* > 0.21, *t*s > 3.04, *p*s_corrected_ < .048), and left hippocampus tracking preceding event memory (*beta* = 0.13, *t* = 3.44, *p*_corrected_ = .025). Finally, to further confirm that our results and conclusions are due to differences in activity related to memory for the upcoming versus preceding trial, and not a fine distinction between offset N and onset N+1, we ran an analysis focused on activity during the 1s ITI, thereby disregarding the offset-onset distinction altogether. The results looked extremely similar to the onset- and offset-based analyses reported above. Thus, we remain agnostic as to the specific time point at an event transition that triggers memory-related activity for the upcoming or preceding event. Full results of all control analyses can be found in our GitHub repository. Therefore, over the course of encoding a multimodal event, there is a temporal transformation in the neural correlates of subsequent memory detail.

### Hippocampal signals integrate episodic features with scene information

To complement the previous onset-related analyses, we next tested the unique influence of subsequent memory for each feature - color, sound, and scene - on brain activity at the end of each event. Whereas there were no significant increases in activity with subsequent color or sound memory, some regions did, however, positively track subsequent scene memory. These effects were not present in the retrosplenial/parahippocampal cortex and occipital regions previously seen; rather, activity of left hippocampus, bilateral lateral prefrontal cortex, and left temporoparietal junction and temporal pole tracked scene encoding at the end of the event (Figure 4A). To visualize the transformation of memory-related activity over encoding, we focused on 5 regions — left hippocampus (L HIPP) and left inferior frontal gyrus (L IFG), selected based on a significant relationship between activity and subsequent memory detail for just-viewed and upcoming event information, respectively, as well as ventral temporal cortex (VTC), auditory cortex (AUD), and retrosplenial/parahippocampal cortex (RSC/PHC), whose onset activity tracked color, sound, and scene encoding, respectively. Here, we predicted memory on trial *N* from two regressors, the standardized activity at both onset *N and* offset *N*, which allowed us to compare their unique effects and control for any low correlation in activity between the beginning and end of a trial (Figure 4B). The relationship between activity and memory detail was significantly greater at event offset than onset in L HIPP (*t* = 2.48, *p* = .016), whereas the opposite was true in L IFG (*t* = −2.42, *p* = .019). AUD and RSC/PHC also showed significantly greater sensitivity to later sound (*t* = −4.82, *p* < .001) and scene memory (*t* = −3.43, *p* = .001), respectively, at event onset than offset, but this pattern was not significant for VTC and color encoding (*t* = −1.13, *p* = .26). Therefore, lateral frontal and visuo-perceptual memory-related onset signals transition to a hippocampal signal after an event that is sensitive to later scene recollection and the amount of information recalled.

**Fig.4.**
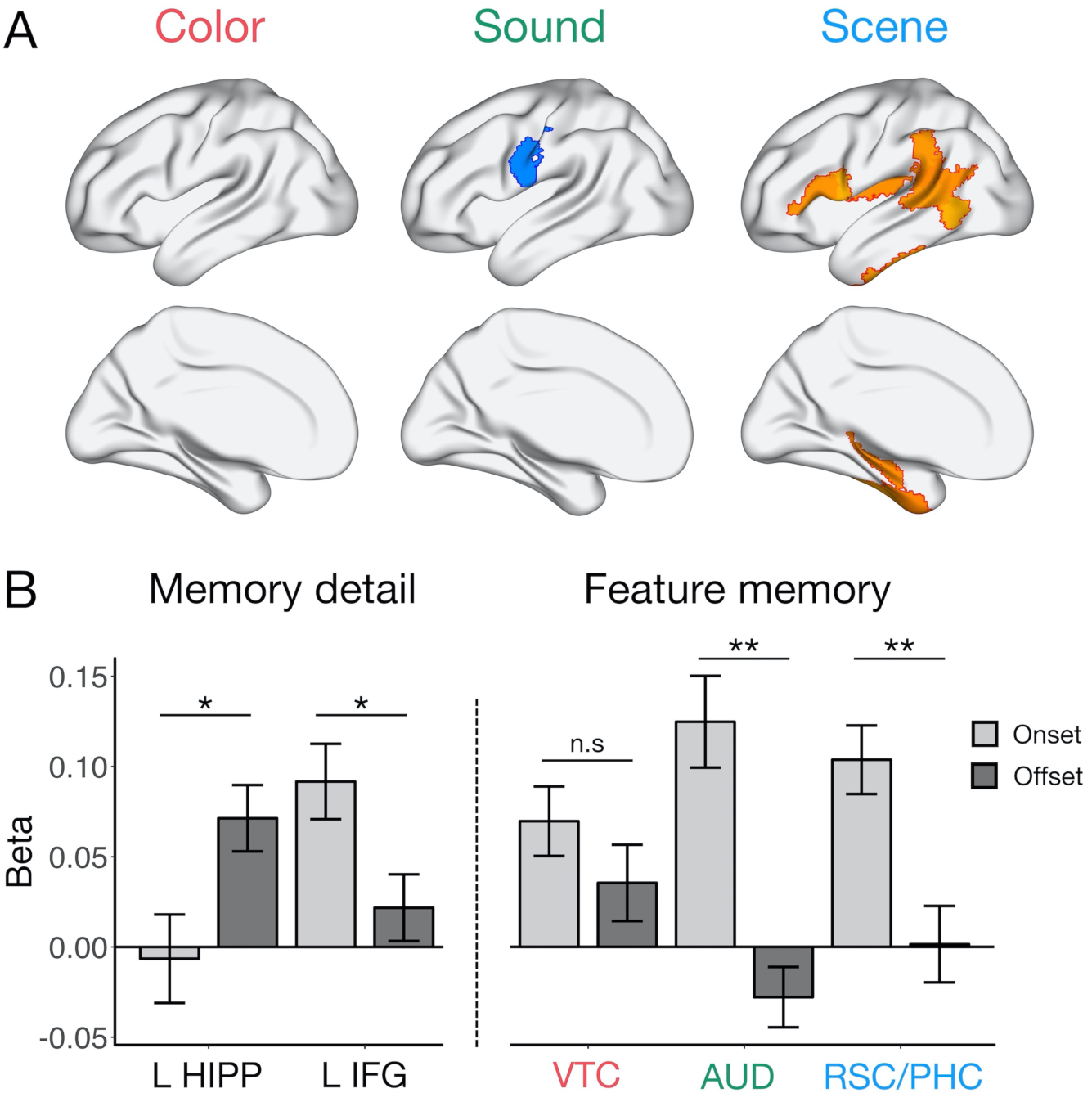
Feature-specific patterns relating activity to memory for the preceding event, and follow-up ROI visualization. A) Feature-specific memory correlates at the offset of each event. Effects shown are significant at *p* < .05 (FDR-corrected). B) Visualization of the change in memory-related activity from the beginning (onset) to end (offset) of each event of interest, for subsequent memory detail (left) in left hippocampus (L HIPP) and left inferior frontal gyrus (L IFG), and for subsequent color, sound, and scene memory (right) in ventral temporal cortex (VTC), auditory cortex (AUD), and retrosplenial/parahippocampal cortex (RSC/PHC), respectively. Bars = fixed effect estimates of memory-related activity, error bars = SEM. ** *p* < .005, * *p* < .05.

The observed pattern of hippocampal activity raises two possibilities about its memory-related function: 1) it may promote memory for the spatial context at an event transition regardless of other episodic features, or 2) it could reflect a binding signal, integrating event features with spatial information after the individual features have been processed. To distinguish these explanations, we predicted the mean activity of left hippocampus with all combinations of features recalled (7 regressors: each feature remembered alone, each possible pair of features remembered together but the third forgotten, or all three features remembered), and tested if hippocampal offset activity is greater in situations where space was remembered with either color or sound features, or both, than when space was recalled in the absence of other information *or* when color and sound were recalled in the absence of spatial information (Figure 5). Indeed, hippocampal activity after an event was only increased (relative to an implicit baseline of no features recalled) when scene information was successfully remembered in conjunction with color, sound, or both features (*t*s > 2.72, *p*s < .009) and not when scene was the only feature subsequently recalled (*t* = 1.00, *p* = .32) or when color and sound were remembered without the associated scene (*t* = 1.26, *p* = .21). Moreover, remembering a conjunction of features that included a scene recruited the hippocampus more than recalling the color and sound without the associated scene (*t* = 1.99, *p* = .046), but the difference between scene alone vs. scene + other feature(s) fixed effects was not significant (*t* = 1.62, *p* = .12).

**Fig.5.**
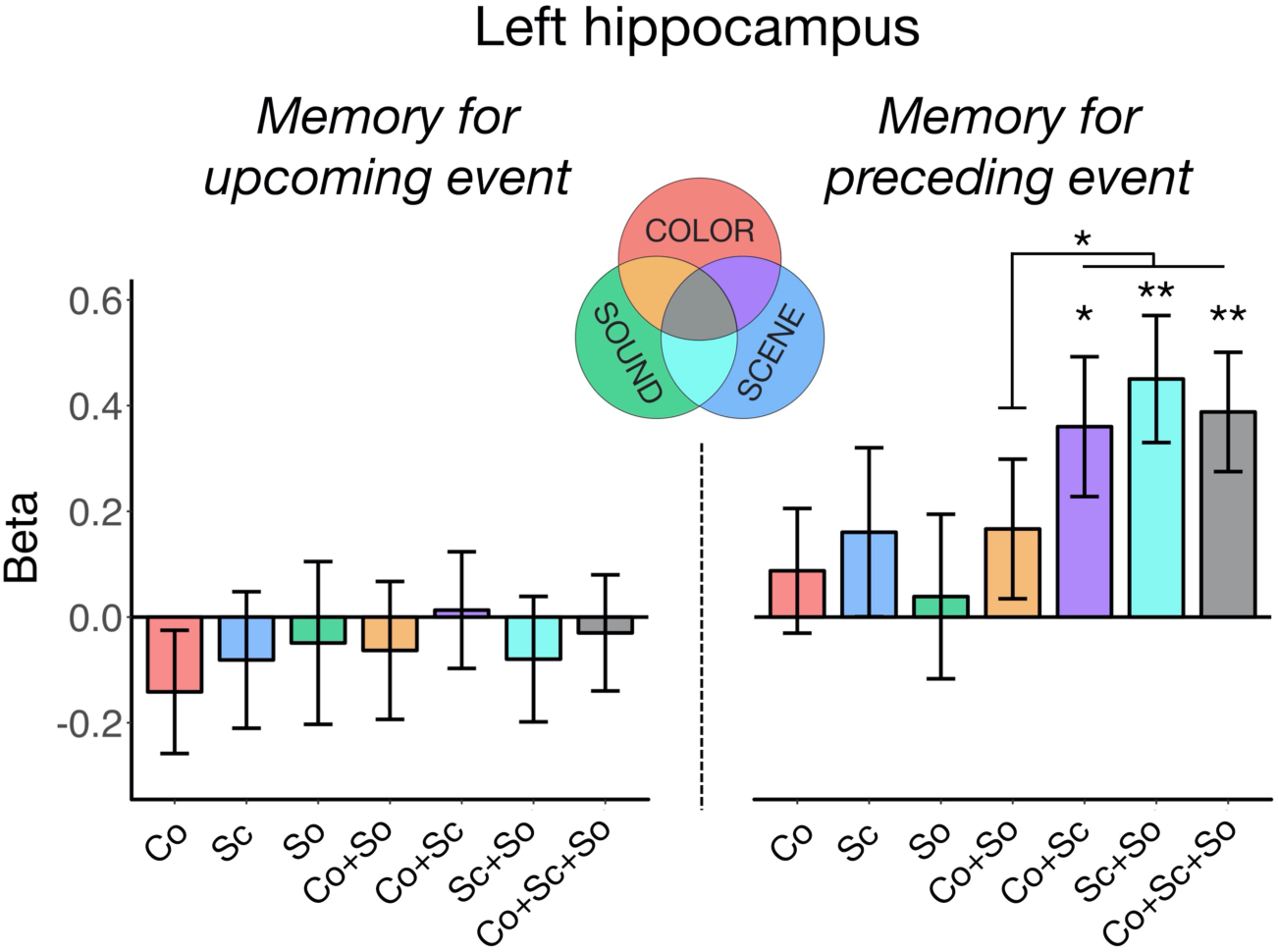
L HIPP activity based on all possible feature conjunctions in memory, relative to an implicit baseline of no features recalled. Hippocampal activity at event onset (left) was not sensitive to memory for any combination of features in the upcoming event. In contrast, hippocampal activity at event offset (right) was increased when spatial information was successfully encoded with other episodic features in the preceding event. Co = Color, Sc = Scene, So = Sound. Bars = fixed effect beta estimates of activity in each condition, error bars = SEM. ** *p* < .005, * *p* < .05.

In an exploratory analysis, we used the same model to determine if this pattern of end-of-event hippocampal encoding activity varied from anterior (head) to posterior (body and tail) segments, determined using the MTL probabilistic atlas (Ritchey et al., 2015). Interestingly, although the activity of both segments increased when more than 1 feature was recalled (anterior: *beta* = 0.25, *t* = 2.58, *p* = .010; posterior: *beta* = 0.23, *t* = 2.41, *p* = .016), selective sensitivity to binding with scenes was a characteristic of only the anterior hippocampus (*beta* = 0.37, *t* = 2.57, *p* = .010; posterior: *beta* = 0.04, *t* = 0.29, *p* = .77). Hence, hippocampal encoding activity, particularly in anterior hippocampus, may act to bind just-viewed features into a coherent spatial event.

## DISCUSSION

This study tested the neural correlates of distinct features encoded in parallel within a complex event, and how those features are integrated into a coherent representation. First, we found that left IFG and visuo-perceptual regions were sensitive to the amount of detail encoded at the onset of each event. Whereas IFG selectively tracked the number of associations recalled, we uncovered unique variance in the brain activity of visuo-perceptual areas predicted by subsequent memory for color, sound, and scene features. In contrast, modeling activity at event offset revealed a shift in the positive neural correlates of successful encoding. Specifically, activity in left hippocampus at the end of each event was predicted by the amount of detail later recalled as well as scene memory. Probing specific feature combinations revealed that these hippocampal effects were driven by binding color and sound information with an associated scene.

Our findings of feature-specific encoding effects demonstrate that multimodal details of complex episodes can be decomposed in the brain, even when encoded simultaneously. The specific pattern of our results converges with univariate and multivariate decoding studies previously using these types of features, including color (Uncapher et al., 2006; Favila et al., 2018), sounds (Gottlieb et al., 2012), and scenes (Park and Chun, 2009; Morgan et al., 2011; Staresina et al., 2011) to study perception and encoding. We additionally showed that these visuo-perceptual neural correlates were only present early on in encoding, which is consistent with relatively earlier subsequent memory effects in ventral visual areas shown using intracranial EEG (Long and Kahana, 2015). Therefore, while the strength of sensory signals predicts upcoming feature-specific encoding, the importance of this sensory activity for memory generally decreases over the course of an event. Moreover, by modeling the unique variance of each feature, we found that scenes showed the most prominent subsequent memory effects. One explanation of this finding is that spatial information serves as the dominant framework for integrating other features within memory (Hassabis and Maguire, 2007; Mullally and Maguire, 2014; Horner et al., 2016; Robin et al., 2016; Bicanski and Burgess, 2018; Robin, 2018). In contrast to many previous studies, memory for spatial information in our task is based on an exact viewpoint rather than overall environment, and it is an open question whether spatial organization extends to precise as well as gist-like contexts. An alternate explanation for these findings is that scenes may be the most salient feature in our task, given that they include a number of other objects and visual details. Thus, despite previous work highlighting the prioritization of spatial information in events, it is difficult to completely disentangle this account from the amount of information contained within spatial contexts.

In addition to feature-specific encoding effects showing an early, transient profile, left IFG also showed an early relationship to total memory detail that diminished over time. This pattern supports the proposal that IFG processes goal-relevant information and initiates an organizational framework (Blumenfeld and Ranganath, 2007; Blumenfeld et al., 2011) to support integration of perceptual features. In our paradigm, participants were asked to generate a story linking the object associations together, and so early IFG activity may help to generate meaningful semantic associations necessary to support feature binding (Gabrieli et al., 1998). Interestingly, and in contrast to prior literature, we did not find a similar relationship between hippocampal onset activity and later encoding success. Instead, we observed a shift whereby left hippocampal activity at the completion of an event significantly predicted the number of just-viewed features recalled. Collectively, these findings suggest that both hippocampus and left IFG contribute to the integration of event components (Staresina and Davachi, 2006; Zeithamova and Preston, 2010) but reveal that their memory signals are temporally dissociable. As such, while left IFG may control organizational strategies, hippocampus may act upon this organizational structure to bind an item with its context. Previous research supports this distinction, showing that IFG generates associations whereas hippocampus is sensitive to an existing relational structure (Addis and McAndrews, 2006), with the former tracking subsequent memory confidence and the latter the number of details remembered (Qin et al., 2011; Mayes et al., 2019). Another interesting contrast suggests that IFG may actively control encoding processes while hippocampal signals are driven by inherent memorability of a scene’s perceptual features (Bainbridge et al., 2017). Furthermore, intracranial EEG work has demonstrated that encoding activity in IFG precedes that in hippocampus (Long and Kahana, 2015), thus providing convincing evidence for their distinct functional and temporal roles in integration during memory encoding.

The absence of hippocampal correlates of subsequent memory early on in encoding is particularly interesting in light of inconsistency in the literature. Although hippocampal encoding effects have often been reported (Spaniol et al., 2009; Kim, 2011), many individual studies have not found them (e.g., Sommer et al., 2005; Haskins et al., 2008; Cooper et al., 2017; Tibon et al., 2019). Although variability in hippocampal encoding effects might be attributed to differences in memory strength and trial selection (Henson, 2005), the current results suggest an additional possible explanation--namely, that such inconsistencies might be related to the duration of the encoding event used in fMRI analyses. Most event-related fMRI tasks use short trials, often 2-3 seconds, and analyze activity locked to the onset, meaning that memory correlates at different times of an event will be virtually indistinguishable. Meanwhile, studies using longer, naturalistic events have consistently shown increased hippocampal activity at the end of an event (Ben-Yakov et al., 2014; Ben-Yakov and Henson, 2018), which correlates with later memory reinstatement both neurally and behaviorally (DuBrow and Davachi, 2016; Baldassano et al., 2017). The hippocampal offset signal is thought to reflect a binding operation based on event discontinuities, linking together all the features that just co-occurred within the same spatial-temporal context before transitioning to a new environment (Staresina and Davachi, 2009), which may be facilitated by rapid reinstatement of the previous event at a boundary (Silva et al., 2019). Evidence that earlier visuo-perceptual activity predicts a hippocampal event boundary response within continuous experience (Baldassano et al., 2017) provides support for this proposal and complements the temporal shift from sensory memory correlates to hippocampal memory effects observed here.

To our knowledge, this is the first study to explicitly test the hypothesis that hippocampal activity at the end of an event reflects binding per se; previous studies have not separately tested memory for distinct kinds of episodic features. Our paradigm allowed us to tease apart which features and conjunctions influenced this signal most, where left hippocampal activity was enhanced selectively on trials were spatial information was successfully remembered *and* integrated with color or sound features, or both. This neural finding corroborates previous explanations of the hippocampal offset effect (Cohen et al., 2015; Ben-Yakov and Henson, 2018) as well as theoretical accounts that hippocampus binds information within a contextual framework (Davachi, 2006; Ranganath, 2010; Eichenbaum, 2017). Moreover, evidence of an end-of-event hippocampal binding signal is also present in other event-related tasks: When learning overlapping associations across trials, hippocampal activity predicts subsequent memory after being exposed to component event associations (Horner et al., 2015) and binds previously studied associations based on shared context (Zeithamova and Preston, 2010; Zeithamova et al., 2012), suggesting a prominent function is the integration of previously encoded event features. While binding appeared to be a common function along the hippocampal long-axis, we additionally found that successful encoding of feature conjunctions involving scenes, specifically, was a more prominent feature of anterior hippocampus, which is frequently associated with scene perception and construction (Zeidman and Maguire, 2016).

When interpreting the results of this study, it is important to consider some limitations of the current approach. Although our events were longer than those in most event-related encoding studies, 6 seconds is still relatively short and as such, we were unable to map out the full time course of memory-related signals over a longer period. This partially reflects the fact that, even though participants were encouraged to generate a story to encode the events, the features were static rather than evolving over time. This design enabled us to measure memory for distinct event features, but future studies would benefit from using longer videos while controlling features that can be tested later on. Relatedly, the lack of jittered event duration means that we cannot definitively assign memory-related effects to the start and end of events. For example, it is possible, yet unlikely, that hippocampal signals have a slower HRF and thus actually reflect an earlier memory-related response. Future studies should therefore use events with variable duration to test the generalizability of the current findings.

In summary, we demonstrated that feature-specific neural responses are dissociable and reflect early encoding processes. Integration of these features into a coherent episode is likely supported by complementary lateral frontal and hippocampal signals, with early left IFG operations possibly providing a meaningful structure, and later hippocampal operations integrating these associations within a contextual framework. This multifaceted process thus promotes subsequent episodic retrieval, helping us to mentally reconstruct the rich detail of our environment.

## ACKNOWLEDGEMENTS

This work was supported by NIH R00MH103401 grant (M.R.). We thank Max Bluestone, Rosalie Samide, and Emily Iannazzi for their assistance with data collection. This research was carried out at the Harvard Center for Brain Science, involving the use of instrumentation supported by the NIH Shared Instrumentation Grant Program; grant number S10OD020039.

## Notes

http://www.thememolab.org/paper-bindingfmri/

